# Analysis of human mitochondrial genome co-occurrence networks of Asian population at varying altitudes

**DOI:** 10.1101/2019.12.21.885905

**Authors:** Rahul K Verma, Alena Kalyakulina, Cristina Giuliani, Pramod Shinde, Ajay Deep Kachhvah, Mikhail Ivanchenko, Sarika Jalan

## Abstract

Networks have been established as an extremely powerful framework to understand and predict the behavior of many large-scale complex systems. We have studied network motifs, the basic structural elements of networks, to describe the possible role of co-occurrence of genomic variations behind high altitude adaptation in the Asian human population. Mitochondrial DNA (mt-DNA) variations have been acclaimed as one of the key players in understanding the biological mechanisms behind adaptation to extreme conditions. To explore the cumulative effects and variations in the mitochondrial genome with the variation in the altitude, we investigated human mt-DNA sequences from the NCBI database at different altitudes under the co-occurrence motifs framework. Analysis of the co-occurrence motifs using similarity clustering revealed a clear distinction between lower and higher altitude regions. In addition, the previously known high altitude markers 3394 and 7697 (which are definitive sites of haplogroup M9a1a1c1b) were found to co-occur within their own gene complexes indicating the impact of intra-genic constraint on co-evolution of nucleotides. Furthermore, an ancestral ‘RSRS50’ variant 10398 was found to co-occur only at higher altitudes supporting the fact that a separate route of colonization at these altitudes might have taken place. Overall, our analysis revealed the presence of co-occurrence interactions specific to high altitude at a whole mitochondrial genome level. This study, combined with the classical haplogroups analysis is useful in understanding the role of co-occurrence of mitochondrial variations in high altitude adaptation.

## Introduction

“I think the next century (21st) will be the century of complexity” once said by Stephen Hawking in context to the wide range of complex systems present around us. To understand and predict behavior of many large scale complex systems, networks provide an extremely powerful framework consisting of nodes and interactions (edges) [1]. For example, complex biochemical activities of a cell can be well understood by underlying protein-protein interaction (PPI) networks. Network framework has been successful in revealing crucial proteins for breast cancer [2], to understand versatility of society [3], to get insights into developmental changes in *C. elegans* [4], and to analyze the progression of biological aging [5]. Motifs, which are complete subgraphs of a network, considered to be building blocks of many complex systems [6]. These motifs have been shown to occur significantly in several biological networks such as gene regulatory networks, ecological networks and neural networks. Two-node motifs have been extensively studied as double negative feedback loops, double positive feedback loops, and auto-activation or repression loops [7, 8].

Motifs are complete subgraphs, and the two node motifs may primarily consist of feed-forward or backward loop. Here we analyzed simple two-node undirected motifs of co-occurring nodes for mtDNA at varying altitudes. The nodes are variable sites and interactions are the co-occurrence of these variations. Co-occurrence of variable sites have been investigated for various diseases, for example in understanding classification and prognosis of acute myeloid leukemia [9], for finding cause of female Duchenne muscular dystrophy [10], for finding co-occurrence of driver mutations in myeloproliferative neoplasms [11]; to understand evolution of influenza viruses [12]; in detection of pesticide resistance in Aedes aegypti [13] and recently in codon level analysis of human mt-DNA which revealed significance of codon-motifs in evolution of human sub-population [14]. The origin and inhabitation of humans in diverse geographical regions across the world has always been a topic of research for anthropologists and geneticists. These studies have pointed out that the presence of environmental diversity in different geographical regions was one of the key factors in causing variability among human groups both at nuclear DNA and mtDNA level [15,16]. A wide range of environmental diversity existed in terms of temperature and altitude driven hypoxia all over the world. One such environment existed in South-Central Asia at Tibetan plateau. The Tibetan plateau was known to be the highest altitude region ever inhabited by humans since the last Largest Glacial Maxima (LGM, 22−18 kya) [17]. The plateau had an average elevation of 4000m above sea level yielding extreme environments such as low oxygen concentration, high UV radiation and arid conditions [18]. The indigenous people of Tibet have acquired an ability to thrive in the hypoxic environment as a result of complex mechanisms of polygenic adaptations (both at nuclear and mtDNA level) [19]. Thus, biological study of the Tibetan plateau was of great interest due to its distinctive environment and migratory profile.

Mitochondria are the energy centers in eukaryotic cells and recent studies showed that the diversity of the mitochondrial genome may have a role in the adaptation to hypoxia in Tibetans [20]. Mitochondria have their own DNA of 16,569 bp encoding 13 proteins and 24 RNAs (2 ribosomal RNAs and 22 transfer RNAs) and are inherited solely through the maternal line (uniparental inheritance). Mitochondria play a regulatory role in oxygen metabolism through oxidative phosphorylation (OXPHOS). Following events may take place during hypoxic exposure; the ATP generation is down-regulated, the activities of respiratory chain complexes are reduced, and reactive oxygen species (ROS) which are produced from the respiratory chain may cause cellular oxidative damage [21-23]. mtDNA mutations that affect OXPHOS could also affect metabolic rate modulation, oxygen utilization, and hypoxia adaptation [24]. Theoretically, it was accepted that migration and genetic drift play crucial roles in controlling mtDNA haplotype frequencies and that mt-DNA variations in a species are selectively neutral [25]. However, recently it was reported that mtDNA variations are the result of natural selection [26, 27]. The factors contributing to these variations are, (i) proteins from mtDNA interact with each other and with those imported from the cytoplasm, and consequently form four of the five complexes of the OXPHOS; and (ii) the presumption of total absence of crossing over in mtDNA, i.e., each genome has a set hierarchical history which is shared by all the genes. Even the highly mutating non-coding control region cannot be assumed to have undergone neutral selection because of indirect genetic effects involving specific loci, affecting mitochondrial transcription and replication in significant ways [28]. Due to these reasons, it was suggested that a site undergoing evolutionary pressure might have equally affected the genealogy of the whole mitochondrial genome [25].

The primary aim of this study was to investigate changes observed in co-occurrence motifs in the human population as altitude is varied, and the secondary being the characterization of these co-occurrence motifs, particularly those depicted changes at genetic levels. We analyzed 673 complete human mitochondrial genomes by categorizing them into eight altitude groups from the sea level up to Tibetan plateau. We kept the interval of 500m between each altitude group taking into account the fact that oxygen percentage decreases by approximately 1% at every 500 meters above sea level [29]. However, on the basis of the elevations, the altitudes below 2000 m are considered as low altitudes while altitudes above 2000 m are considered to be moderate to high altitudes [30]. We have constructed two types of the networks, one by considering co-occurring variants as nodes and another by considering the genes consisting of these co-occurring variants as nodes (Fig 1). Formal Random Forest classification based on variable sites as features demonstrated that the non-adjacent groups can be discriminated with at least 80% accuracy for the test set with 5-fold cross validation. The co-mutational cohorts were defined in terms of complete subgraphs aka motifs. The two nodes motifs considered in this paper consisted of variable sites in which minor alleles co-occurred equal to or above the set threshold of the co-occurrence frequency (C_th_). These co-occurrence motifs revealed that the variable sites co-occurred within their own gene or gene complexes which are formed by multiple genes, emphasizing intra-genic constraint of genomic evolution. In addition, similarity analysis and phylogenetic distance based Principal Coordinates analysis bifurcated all the altitude groups into two categories, lower and higher altitude regions. A characterization of co-occurrence motifs at genetic level revealed the dominant role of cytochrome b in the adaptation at Tibetan plateau. Additionally, we analyzed the deleteriousness of selected variants using CADD scores and HmtVar predictions.

**Figure 1.** (a) Total variable sites are extracted from a particular group and co-occurrence threshold is applied (See Methods, Section 2.2). (b) For each sample, one set of motifs were constructed. (c) These motifs were then merged to construct one master network where nodes were variable sites. (d) This master network was then used to construct a gene-gene interaction network by mapping the variable sites corresponding to each gene.

## Results

### Characterization of Variable sites

A total of 3,829 variable sites exist for all the altitude groups. Out of which 3127 variable sites took part in mutation cohort formation of motifs of order two or higher. However, here we have considered the simplest motifs of order two only for the further analysis. Among these variable sites, □65% sites were found to be located in protein-coding regions (overlapped sites were double counted) with the rest lying in non-coding regions (control region, t-RNAs and r-RNAs) (Table 1). This was not surprising as 11,395 (□68%) sites out of the total 16,569 sites belong to the coding region. Most of the variable sites were bi-allelic, a few were having three alleles (tri-allelic) in all the groups and only group 3 had one site with four alleles (Table 1). These sites are well documented in the Mitomap database for various genomic studies. Although much about the tri-allelic sites have not been understood, nonetheless their presence has been shown to be responsible for natural selection [31]. Approximately 90% of the variable sites were transitions, yielding a high transition to transversion ratio (Ts/Tv) (Table 1) which has already been reported [32] to be responsible for the conservation of structures at protein level among the individuals within a species [33]. In the context of varying altitudes, this ratio remained high which further emphasized the importance of survival and for mitochondrial functionality. As we know that the functional stability comes from the structural integrity of proteins and the structural integrity arises due to specific interactions of amino acids [34]. These interactions of amino acids, in turn, were shown to be affected by mutations and their co-occurrence in the genome [35].

**Table 1:** Statistics of Variable sites

| Altitude group | Variable sites | Coding region sites | Non-coding region sites | Ts:Tv | No. of Tri-allelic sites |
| --- | --- | --- | --- | --- | --- |
| Group 1 | 474 | 286 (60.33\%) | 188 (39.6\%) | 16.5:1 | 5 |
| Group 2 | 435 | 270 (62.06\%) | 165 (37.94\%) | 14.6:1 | 6 |
| Group 3 | 644 | 423 (65.68\%) | 221 (34.32\%) | 14.2:1 | 7 |
| Group 4 | 694 | 432 (62.24\%) | 262 (37.76\%) | 12.5:1 | 11 |
| Group 5 | 429 | 274 (63.86\%) | 155 (36.14\%) | 17.8:1 | 4 |
| Group 6 | 354 | 222 (62.71\%) | 132 (37.29\%) | 22.7:1 | 1 |
| Group 7 | 357 | 212 (59.38\%) | 145 (40.62\%) | 17.0:1 | 1 |
| Group 8 | 442 | 279 (63.12\%) | 163 (26.88\%) | 13.3:1 | 1 |

### Discriminating Altitude Groups

According to the dataset, there was no unique correspondence between haplogroups and altitude groups. Therefore, we investigated the degree of difference between altitude groups referring to the whole set of mt-DNA variable sites. This was achieved by applying several methods of classical genetic analysis and machine learning approaches. First, we built the phylogenetic tree for the sequences with excluded positions that are defining haplogroups. For the phylogenetic distances between all subjects’ pairs, we applied PcoA and plot 2 first Principal Coordinates (Figure 2). This analysis revealed the existence of two main clusters, one cluster with individuals living above 2000 m and another cluster with the individuals living below 2000 m of altitude. However, this method works only with the whole sequence and does not allow to highlight single variants.

**Figure 2:** The 2 main Principal Components for the phylogenetic distances. Each point represents one subject, its color corresponds to the altitude group.

Next, we applied the formal machine learning Random Forest binary classification for each group pair, treating variable mt-DNA sites as features. The results demonstrated a good classification accuracy of over 80% for the pairs of groups having altitude different between them greater than 2000m (Table 2), thus confirming the altitude association of the studied mt-DNA samples. However, the output lists of SNPs ranked by importance did not inform whether the definitive mutations are independent or co-occurring. This would be the general case for machine learning models, while the following co-occurrence network analysis allowed us to go beyond.

**Table 2:**
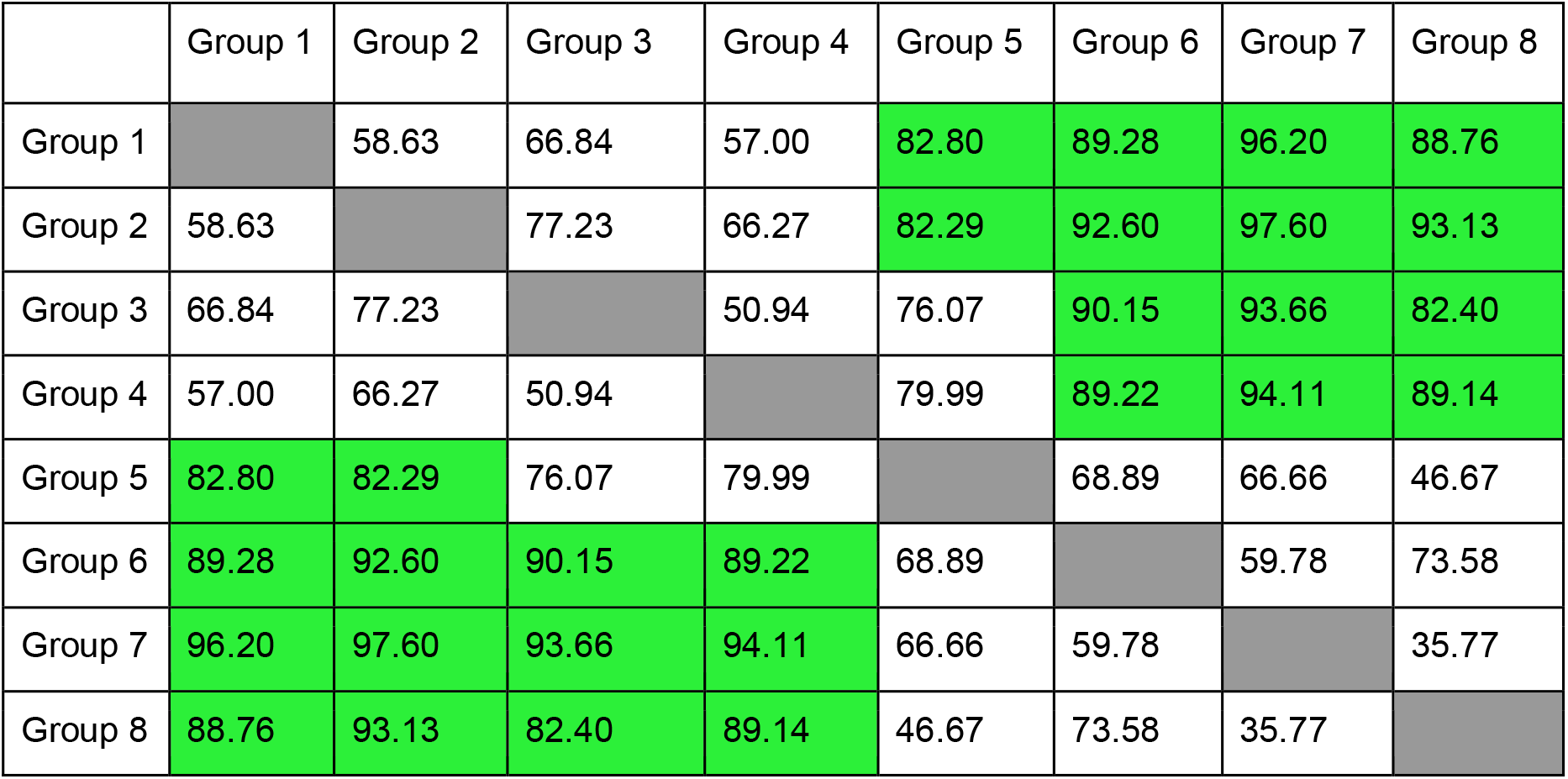
Accuracy of binary classification between altitude groups, the cases of good discrimination, over 80%, are highlighted by green

The classical genetic analysis methods reveal certain mt-DNA regions which help to distinguish subjects living lower and higher than 2000 m. DAPC (Discriminant analysis of Principal Components) shows good differentiation between subjects belonging to different altitudes for the D-loop (Control region), MT-ATP6, MT-ND1, MT-ND2, MT-ND4, MT-ND5 (examples of regions with and without clustering are on Figure 3). The pairwise Fst statistic shows good differentiation between subjects belonging to different altitudes (with p-values < 0.05 for all group pairs with altitude difference < 2000m) for the following regions: D-loop (Control region), MT-RNR1, MT-RNR2, MT-ATP6, MT-CO1, MT-CO2, MT-CO3, MT-CYB, MT-ND1, MT-ND2, MT-ND4, MT-ND5, MT-ND6.

**Figure 3:** The 2 main Principal Coordinates, obtained by DAPC, for the D-loop (left) and ATP8 (right). Each point represents one subject, its color corresponds to the altitude group. For D-loop there are 3 main clusters, corresponding to altitudes 0-2000m, 2501-3000m, 2001-2500 and 3000-4001m. For the ATP8 there are no separate clusters.

DAPC and Fst exhibited similar results for D-loop (Control region), MT-ND1, MT-ND2, MT-ND4, MT-ND5, showing good differentiation between those pairs of the groups having altitude difference more than 2000m. The main limitation of these methods was that they worked with sequences or certain regions and did not reveal any significant mutated positions and co-occurring ones.

### Categorization of Variable sites

We categorized variable sites based on their occurrence in the network as follows; (i) Isolated nodes (variable sites which did not take part in network construction) and (ii) Connected nodes (variable sites which took part in network construction). Further, these two types of nodes were sub-categorized into (a) Global nodes (the nodes present in all the altitude groups), (b) Local nodes (the nodes presented exclusively in a particular group) and (c) Mixed nodes (the nodes presented in more than one altitude groups but not in all). This categorization helped us to decipher the role of variable sites in terms of two nodes co-occurrence motifs. It was also observed that the percentage of local connected nodes was decreasing with increasing altitude while the percentage of local isolated nodes was increasing with increasing altitude. Further, the number of links (Nc) provided the information about the co-occurrence pairs formed by the connected nodes having perfect co-occurrence frequency. The average degree (<k >) (the average number of connections of each node) of the network was found to be nearly similar in all the networks and also confirmed sparseness of the networks of all the groups. The real-world complex networks are found to be mostly sparse [36]. Here, the sparsity of co-occurrence networks meant that the variable sites are having very few perfectly co-occurring pairs. The same are summarized in Table 3.

**Table 3:** Categorization of Variable sites (nodes)

| Altitude groups | Connected Nodes | | | Isolated Nodes | | | $N_c$ | $\langle k \rangle$ |
| --- | --- | --- | --- | --- | --- | --- | --- | --- |
|  | Local (%) | Mixed (%) | Global (%) | Local (%) | Mixed (%) | Global (%) |  |  |
| Group 1 | 31.5 | 67.5 | 1 | 28.4 | 16.2 | 55.4 | 1855 | 9 |
| Group 2 | 29.4 | 69.5 | 1.11 | 29.7 | 14.9 | 55.4 | 1324 | 7 |
| Group 3 | 36.9 | 62.3 | 0.77 | 32.3 | 34.7 | 33 | 1658 | 6 |
| Group 4 | 38.2 | 61.1 | 0.72 | 37.5 | 32.3 | 30.2 | 2219 | 8 |
| Group 5 | 23.3 | 75.6 | 1.15 | 32.1 | 17.3 | 50.6 | 1443 | 8 |
| Group 6 | 20.4 | 77.9 | 1.34 | 25.5 | 14.5 | 60 | 1182 | 8 |
| Group 7 | 27.1 | 71.5 | 1.44 | 37.5 | 11.3 | 51.2 | 950 | 7 |
| Group 8 | 25.9 | 73.1 | 1.1 | 35.9 | 11.5 | 52.6 | 1409 | 8 |

Isolated Nodes: A total of 702 isolated nodes were found in all the groups with an average of 88 nodes per altitude group (□25%) for the threshold considered (C_th_ = 1) here. The isolated nodes represent molecular sites which undergo independent variation in the context of motif formation as these variable sites are not co-occurring with any other variations. To gain further insights into these nodes, we mapped them with their corresponding genes and found that a significantly high number (□50%) of these nodes belonged to the control region (CR) in each altitude group (Fig 4). This significantly high contribution of CR in the isolated nodes suggests that the variants in this region are independent of the variants in other regions. It is already known that CR undergoes a higher rate of mutations as compared to the other mitochondrial genes [37] and that CR is characterized by hypervariable regions (HVR) that are mutational hotspots. These isolated nodes also contain the variable sites which showed a high minor allele frequency (MAF) of nearly 40%. If the frequency of a minor allele in any given population is high, it is considered that the major allele is far from being fixed and instead contributes to the genetic diversity in that particular population. The presence of variable sites with such a high MAF in the isolated nodes and their correspondence to CR again suggest that this region has undergone a higher rate of mutation as compared to other (non)coding regions in mt-DNA. Further, all the protein-coding genes and RNA genes contributed less than 10% to the isolated nodes at all the altitudes. Among the protein-coding genes, ND5 showed the highest percentage followed by MT-ND2, MT-CYB, MT-ND4, MT-CO1 and MT-ATP6 while MT-ATP8 showed the lowest contribution in the isolated nodes (Fig 4).

**Figure 4:** The mean of the number of isolated nodes for corresponding genes across all the altitudes.

Global Nodes: 41 variable sites in the isolated nodes and only 4 variable sites from the connected nodes were found to be common in all the altitude groups, which were referred as ‘global isolated nodes’ and ‘global connected nodes’, respectively. It was quite apparent that the global connected nodes were found to be 10-fold less than the global isolated nodes. This means that the variations which were occurring globally were not necessarily co-occurring perfectly with the other variable sites and their co-occurrence might be population specific or more precisely in our case, altitude specific. The four global connected nodes were 9540, 10400, 10873 and 16327. Out of these four nodes, three, 9540, 10400 and 10873 were found to be connected with each other forming one single motif of order three in all the altitude groups except the group 3 where only 9540 and 10873 formed a motif whereas in group 4, despite having the maximum number of nodes, surprisingly not a single node co-occurred with these three global connected nodes. The 16327 node formed a star-like network structure with the other nodes in all the groups. This star-like topology suggested that 16327 is acting as a root node for expansion of this haplogroup marker [38]. It is well known that 16327 is a major haplogroup marker of M haplogroup, a haplogroup specific to Asian population. Further, 9540, 10400, 10873 and 16327 belonged to MT-CO3, MT-ND3, MT-ND4 and CR genes, respectively. The genes corresponding to nodes which were co-occurring with global nodes of the coding region (9540, 10400 and 10873) are summarized in Table 4. From the table it is quite evident that these three global nodes preferred to co-occurred with the nodes belonging to the coding regions. The marker 10398 is an ancestral one belonging to L3 lineage, universally present in M haplogroup and found to co-occur with global nodes present only in higher altitude groups 6 and 7.

**Table 4:** Genes corresponding to variable sites co-mutating with global connected nodes of coding region

| Altitude group | Genes |
| --- | --- |
| Group 1 | MT-ATP6 (8701), MT-CYB (14783), MT-CYB (15301) |
| Group 2 | MT-ATP6 (8701), MT-CYB (15043), MT-CYB (15301) |
| Group 3 | MT-ATP6 (8701), MT-CYB (14783), MT-CYB (15301) |
| Group 4 | - |
| Group 5 | MT-ATP6 (8701), MT-CYB (14783), MT-CYB (15043), MT-CYB (15301) |
| Group 6 | MT-ATP6 (8701), MT-ND3 (10398), MT-CYB (14783), MT-CYB (15043), MT-CYB (15301) |
| Group 7 | MT-ATP6 (8701), MT-ND3 (10398), MT-CYB (14783), MT-CYB (15043), MT-CYB (15301) |
| Group 8 | MT-ATP6 (8701), MT-CYB (14783), MT-CYB (15043), MT-CYB (15301) |

Local Nodes: The variable sites which co-occurred exclusively in a particular group were considered as local nodes. Using a two-proportion Z-test, we found that in the higher altitude groups, the local connected nodes decreased significantly as compared to that of the lower altitude groups. To find the difference at the gene level between the local connected nodes of each pair of the altitudes, we mapped local nodes to genes (Fig 5). Mapping with genes revealed a very peculiar property of the mitochondrial genome. Although the local nodes were exclusively mutated sites for a particular altitude which were not found in any other altitude regions, at the genetic level the variations remained similar for all the altitude groups. This is because different variations might belong to the same gene. Among all the gene complexes, ND genes showed the highest count which was expected as ND complex comprises seven sub-units. Apart from that we looked for co-occurrence pairs formed by these local nodes only. We found that the lower altitude groups (group 1 to group 4) exhibited a very high percentage of such pairs than the higher altitude groups (group 5 to group 8) as well as the corresponding random networks.

**Figure 5:** (a)The local nodes were mapped to genes and gene counts were plotted. Each gene was showing a similar count in each altitude. (b) The percentage of co-occurrence pairs consisting only of local nodes.

High altitude markers: The variable sites 3010, 3394 and 7697 were reported as high-altitude markers and have been associated with high altitude adaptation in Tibetan population [20]. The markers 3010, 3394 and 7697 belonged to MT-RNR2, MT-ND1 and MT-CO1 genes, respectively. Further, we wanted to investigate the motifs derived by these markers to determine their co-occurrence background. The marker 3010 was found only in groups 1, 6, 7 and 8. In groups 6, 7 and 8, it formed the motif with 14668 (MT-ND6) and 8414 (MT-ATP8) whereas in group 1 it formed a separate cohort with 295 (Control region), 462 (Control region), 8269 (MT-CO2), 12612 (MT-ND5) and 16069 (Control region). The marker 3394 was found in group 1 to form motif with 11335 (MT-ND4), in group 4 to form motif with 4832 (MT-ND2) and 16258 (CR) and in groups 5 and 8 to form motifs with 1041 (MT-RNR1), 4491 (MT-ND2) and 14308 (MT-ND6) in both the groups. The marker 7697 was found only in higher altitude groups 5, 6, 7 and It formed motifs with 711 (MT-RNR1), 7142 (MT-CO1), 9242 (MT-CO3) in group 5, with 453(CR), 711 (MT-RNR1), 7142 (MT-CO1), 9242 (MT-CO3) and 14417 (MT-ND6) in group 6, 711 (MT-RNR1), with 7142 (MT-CO1), 9242 (MT-CO3) and with 14417 (MT-ND6) in groups 7 and 8. Further, our analysis revealed that these high altitude markers tended to co-occur with their own gene complexes which emphasized the intragenic influence on co-occurrence of nucleotides.

### Grouping of altitudes based on co-occurrence motifs

Jaccard similarity coefficient was used to find out the similarity between each altitude group using the mixed nodes (Fig 6). This similarity coefficient led to the distinction of two major clusters, one with groups 1, 2, 3 and 4 (lower altitude cohort) and the other with groups 5, 6, 7 and 8 (higher altitude cohort). The nodes of the dendrogram in Fig. 6 represented altitude groups. Two altitude groups were found to form doubletons within each cluster. Moreover, it was deduced from the dendrogram that the human population splits up into two subpopulations giving rise to lower altitude cohort and higher altitude cohort. The lower altitude cohort further segregated into two sub-groups forming one clade with Grp 2 and Grp 3 and another clade with the Grp 1 and Grp 4. The higher altitude cohort further segregated into two sub-groups forming one clade with the Grp 5 and Grp 8 and another clade with the Grp 6 and Grp 7. It is noteworthy that in the lower altitude groups, Grp 1 and Grp 4 descended from a common sub-group. Similarly, in the higher altitude groups, Grp 5 and Grp 8 descended from a common sub-group despite having geographical distances between these altitude ranges. Many previous studies have pointed out that early humans have migrated towards higher altitudes in summer for hunting, whereas moved towards lower altitudes during winter season to avoid extreme harsh environments [39]. This seasonal migration is common even today in the plateau [40]. The segregation of the human population in lower and higher altitude groups observed in our analysis suggests that humans may have migrated through discrete pathways searching for a better environment for establishment.

**Figure 6:** Cluster dendrogram was produced using common nodes between each altitude. It is clearly observed that two clusters are formed, one with groups 1 to 4 (lowest to middle) and other with groups 5 to 8 (middle to highest).

### Gene-gene interactions

Gene-gene interaction networks were compared with the corresponding random networks. We found that only 72 (□10%) of all possible gene-pairs were significantly deviated (falling out of the standard deviation range) from random networks (Fig 7). Moreover, out of total 72 gene-pairs, 46 were found to be present in any one of the altitude groups, 23 were found to be present in any two of the altitude groups, and only 1 pair was found to be present in any 3, any 4 and any 7 altitude groups. There were certain pairs in which one of the genes was tRNA such as in group 1 tRNA-Gly. The gene-pair CYB-CR was found to be present in all the groups except in the 5th group where these genes were present but interacted with other genes. Further, this pair had more weight than that of the corresponding random network in group 6 while less weight than that of the corresponding random networks in the other groups.

**Figure 7:** Gene-gene interaction pairs which showed deviation from random networks.

Interestingly, there existed only three genes which formed pairs with themselves, these genes are MT-CYB, MT-ND5 and CR. The gene-pair CYB-CYB was found to be present in the groups 4 and 8. In group 4, its weight was found to be less than that of the corresponding random network, while in group 8 its weight was found to be more than the corresponding random network. The other gene-pair CR-CR was also found in group 2 and group 8. Moreover, in group 8, in seven out of fifteen gene-pairs one of the genes was MT-CYB. Another gene-pair CO3-ND6 was found to be present in groups 6 and 7. Apart from the CR and the coding genes, despite having small length and less variable sites, certain tRNA genes were also found to form gene interaction pairs (Table 5). Particularly tRNA-Gly was present in groups 1, 2, 3 and 8. Interestingly, tRNA-Thr exhibited the highest number of variable sites among tRNA genes, but it was found to be present only in group 3 co-occurring with MT-CO1 gene. tRNA-Met was present only in groups 5 and 8. Overall, we found different gene-gene interactions at varying altitudes which can be further analyzed for possible adaptation or disease association.

**Table 5:** The number of genes and genes pairs in each altitude group along with the specific tRNA genes

| Altitude Group | No. of Gene-pairs | No.of Gene-pairs | tRNA genes |
| --- | --- | --- | --- |
| Group 1 | 14 | 13 | Gly, Phe |
| Group 2 | 7 | 7 | Gly |
| Group 3 | 12 | 15 | Ala, Asp, Gly, Thr |
| Group 4 | 30 | 16 | His, Ser |
| Group 5 | 10 | 15 | Arg, Met |
| Group 6 | 9 | 10 | --- |
| Group 7 | 9 | 13 | Glu |
| Group 8 | 15 | 14 | Gly, Met |

### Impact of Variants

To explore the predicted functional impact of selected variants, we extracted the various prediction scores from HmtVar database [41] and Combined Annotated Depletion Dependent (CADD) database [42] (Table 6). The table shows the prediction scores for various variants which were found to be significant based on the co-occurrence networks. The C-scores and PhyloP scores conveyed about the deleteriousness and conservation score, respectively. The positive PhyloP score predicted conserved sites while its negative value predicted fast-evolving sites. For most of the variants HmtVar scores were unavailable while we discussed the relative significance of those which were available. The high-altitude markers were shown to have only polymorphic nature with some degree of pathogenicity. The variant T3394C was predicted to have high pathogenicity along with disease-causing impact by MutPred, PhD SNP and SNPs & GO databases. This variant has been associated with Leber’s hereditary optic neuropathy (LHON), diabetes mellitus, osteoarthritis, cardiomyopathy in non-Asian populations while in Asian population it has been associated with high-altitude adaptation. The co-mutating partners of T3394C were all fast-evolving sites except G4491A. There are four variants which co-occurred with G7697A in which two variants were highly evolving while two variants were found to be conserved. For most of the variants the predictions were not available in HmtVar while PhyloP score was available for all the variants. The PhyloP scores of pairs of Global connected nodes suggested that the Global connected nodes tend to pair with conserved sites across all the altitude groups.

**Table 6:** HmtVar predictions and CADD scores for selected variants

| S. No | Variant | AA | HmtVar Prediction | Mut Pred | PP HumDiv | PP HumVar | PhD SNP | SNPs & GO | PhyloP | C-score |
| --- | --- | --- | --- | --- | --- | --- | --- | --- | --- | --- |
| **High Altitude Markers** |  |  |  |  |  |  |  |  |  |  |
| 1 | C3310T | P→S | Likely Polymorphic | Low Path | Benign | Benign | Neutral | Neutral | -0.1 | 2.5 |
| 2 | T3394C | Y→S | Polymorphic | High Path | Benign | Benign | Disease | Disease | 7.0 | 1.5 |
| 3 | G7697A | V→I | Polymorphic | Low Path | Benign | Benign | Neutral | Disease | 7.8 | 0.4 |
| **Global Connected Nodes** |  |  |  |  |  |  |  |  |  |  |
| 4 | T9540C | Syn | NA | NA | NA | NA | NA | NA | 9.6 | 0.3 |
| 5 | C10400T | T→A | NA | NA | NA | NA | NA | NA | 5.8 | 1.3 |
| 6 | T10873C | Syn | NA | NA | NA | NA | NA | NA | -1.4 | 0 |
| 7 | C16327A | Reg | NA | NA | NA | NA | NA | NA | -2.5 | 1.1 |
| 8 | C16327T | Reg | NA | NA | NA | NA | NA | NA | 0.11 | 1 |
| **Global Connected Nodes Pairs** |  |  |  |  |  |  |  |  |  |  |
| 9 | A8701G | T→A | Polymorphic | Low Path | Benign | Benign | Neutral | Neutral | 2.0 | -0.7 |
| 10 | A10398G | T→A | Polymorphic (Protective) | Low Path | Benign | Benign | Neutral | Neutral | 7.2 | -1 |
| 11 | T14783C | Syn | Likely Pathogenic | NA | NA | NA | NA | NA | 0.11 | 0.3 |
| 12 | G15043A | Syn | Likely Pathogenic | NA | NA | NA | NA | NA | 5.26 | 0.7 |
| 13 | G15301A | Syn | Likely Pathogenic | NA | NA | NA | NA | NA | 7.5 | 0.7 |
| **Pairs of 3010** |  |  |  |  |  |  |  |  |  |  |
| 14 | C14668T | Syn | NA | NA | NA | NA | NA | NA | 3.62 | 1 |
| 15 | C8414T | L→F | Likely Pathogenic | Low Path | Damaging | Damaging | Disease | Neutral | 2.61 | 2.4 |
| 16 | C295T | Reg | NA | NA | NA | NA | NA | NA | -0.07 | 1 |
| 17 | C462T | Reg | NA | NA | NA | NA | NA | NA | 1.0 | 1.1 |
| 18 | G8269A | Syn | NA | NA | NA | NA | NA | NA | -6.4 | 1.1 |
| 19 | A12612G | Syn | NA | NA | NA | NA | NA | NA | 0.12 | 0.6 |
| 20 | C16069T | Reg | NA | NA | NA | NA | NA | NA | -4.0 | 1 |
| **Pairs of 3394** |  |  |  |  |  |  |  |  |  |  |
| 21 | A166258T | Reg | NA | NA | NA | NA | NA | NA | -1.4 | 1 |
| 22 | A1041G | Reg | Likely Benign | NA | NA | NA | NA | NA | -4.0 | 0.8 |
| 23 | G4491A | V→I | Polymorphic | Low Path | Benign | Benign | Neutral | Neutral | 0.14 | -1.2 |
| 24 | T14308C | Syn | NA | NA | NA | NA | NA | NA | -0.8 | 0.2 |
| 25 | T453C | Reg | NA | NA | NA | NA | NA | NA | -4.2 | 1 |
| **Pairs of 7697** |  |  |  |  |  |  |  |  |  |  |
| 26 | T711C | Reg | NA | NA | NA | NA | NA | NA | 0.2 | 0.8 |
| 27 | T7142C | Syn | NA | NA | NA | NA | NA | NA | -20 | -0.7 |
| 28 | A9242G | Syn | NA | NA | NA | NA | NA | NA | -11.4 | 0.2 |
| 29 | A14417G | V→A | Likely Polymorphic | Low Path | Benign | Benign | Neutral | Neutral | 1.0 | 0.3 |

The C-scores for all the nodes for each altitude-group are shown in Figure 8. The C-scores are ranging between −2 to 4. Although the absolute C-scores had no meaning, they did have a relative meaning. The negative value predicted a site to be proxy neutral while a positive value predicted it to be a proxy deleterious site. It was observed that C-scores of the variants from the control region were close to 1 which suggested that the variants in the control region were neither deleterious nor beneficial across varying elevations. The C-scores for selected variants were mentioned in Table 6 where the highest deleterious score was 2.5 for C3310T variable site, however, this variant was exhibited to have low pathogenicity and likely to be neutral. By projecting the C-scores for all the nodes for each master co-occurrence network, we were able to look at the likely role played by each variant in contributing proxy-deleterious or proxy-neutral variants for co-occurrence network construction. We found that more than 60% nodes were falling in between 0 to 1 C-scores, ∼30% were found to show >1 C-score and ∼10% were found to show <0 C-scores. This suggested that a few individual variants might have deleterious effects on population, however, mutational cohorts might be able to subsidise these predicted deleterious effects.

**Figure 8:** The CADD scores are plotted for all the nodes of each master co-occurrence network. The negative values show proxy neutral predictions while the positive values show proxy deleterious predictions for each variable site.

## Discussions and Conclusion

We investigated altitude driven co-occurrence of variations in Tibetan and lower altitude population using the two nodes network motifs. Phylogenetic techniques, as well as classical population genetic analysis methods had been applied. As such, DAPC, Fst and machine learning approaches allowed us to build formal rules to discriminate different altitude groups with good accuracy but failed to distinguish between the independent and co-occurring mutations. Here, we had used a network model to represent the mitochondrial genome as a complex genomic interaction network based on the co-occurring nucleotide pairs over the entire genome. Even though we took a perfect co-occurrence frequency, nearly 75% nodes (variable sites) took part in the network construction suggesting a widespread presence of mutual variations in the human mitochondrial genome. This widespread of mutual variations further suggested that these variations richly co-occur with each other. The rest of the 25% nodes, which we categorized as the Isolated nodes, were mostly contributed by the Control region which was a mutational hotspot in human mtDNA. These isolated nodes were also found to be more in the population belonging to the lower altitude groups as compared to those at the higher altitude groups. An absence of any selective pressure in lower altitude groups might be a reason for lower co-occurrence of the variable sites yielding high number of isolated nodes (more independent signals). Whereas, for the high-altitude groups which were accompanied with peculiar conditions (oxygen, temperature, UV, etc) leading to more selective pressures which might be causing co-occurrence of the variants for their positive advantages. Further studies are needed to correlate these observations with peculiar phenotypes.

Another category of the nodes, the local nodes, constituted nearly 30% of total nodes for a particular group. The presence of local nodes provided evidence that the human population had a specific signature at nucleotide variation level for varying altitudes, which might be arising by different genetic history of the population that colonized a particular area especially in the low/medium altitudes where the intensity of natural selection on human mt genome was likely to be low. This signature seemed to disappear when we mapped these local nodes to their corresponding genes and counted the number of local nodes for each gene complex. This loss of the signature revealed a peculiar property of the mitochondrial genome that even with exclusive variations, the genomic functionality remained undisturbed, keeping fundamental molecular functions intact. This did not imply that the environment did not affect the adaptive function of the mitochondrial genome but that the neutral variation was a common and well described phenomenon for the mitochondrial genome. Further, the co-occurrence analysis of the high-altitude markers revealed the presence of intra-genic constraint in the high-altitude population. This suggested that the presence of particular variation was not sufficient for adaptation, but that variation had to be assisted by other variations in the same gene or same gene complex. Here we had analyzed only mt-DNA but it was likely that co-occurrence occurred between mt-DNA and the nuclear genomes as the role of nuclear variants in adaptation to high altitudes was well described [43, 44]. Further studies are necessary to identify these interactions between the two genomes. Variable site 711 of 12S rRNA gene was found to co-occur with both the markers 3394 and 7697. The variants 711C and 14417G defined the sub-haplogroups M9a1a1c1b1a, M9a1a1c1b1a1 and M9a1a1c1b1a2 which were widely distributed in East Asian and Southeast Asian populations, and prevalent in Tibetan population. Moreover, MT-RNR1 gene encoded for a protein responsible for regulating insulin sensitivity and metabolic homeostasis [45]. Particularly, the co-occurrence of 7697 with 14417 was observed in the group 7 and 8. Since, the biological effects of high altitude were observed at >3000 m existence of these co-occurrence pairs seemed a possible combination of variants that affects mitochondrial bioenergetics. Variant 4491 of ND2 gene was found to be associated with high altitude pulmonary edema (HAPE) susceptible in low altitude population [46] whereas in our analysis, this variant was found to co-occur with 3394 and its exclusive presence in the higher altitude regions suggested its adaptive dependence. The variable sites forming the global connected nodes 9540 and 10873 were the ‘RSRS50’ ancestral variants [47] while 10400 and 16327 were markers of M sub-haplogroups [46]. The markers 9540 and 10873 were found to be present throughout the human population, 10400 was known to be specific to Asian population and 16327 was known to be the marker of C sub-haplogroup of M haplogroup which was specific to Siberian and American regions. The presence of 16327 in Asian population was not surprising since human migration in American continent took place through the Beringia bridge from Siberia [49,50]. Phylogeny based study had shown that 10398 resulted through selective sweep at colder geographical regions [51] which was supposed to lower the oxidative phosphorylation coupling leading to release of more heat. This possible tradeoff between ATP generation and thermogenesis seemed to play a key role in adaptation at colder higher altitudes. Substitution from A to G at 10398 corresponded to substitution of an alanine amino acid residue by a threonine at the carboxyl end of ND3 gene, a subunit of NADH-ubiquinone oxidoreductase (complex I). The presence of 10398 exclusively at higher altitudes suggested that a separate ancestral population might have colonized these regions. We found the existence of three variable sites belonging to the MT-CYB gene which were co-occurring with global connected nodes; these variable sites are 14783, 15043 and 15301. Although we had collected the mt-DNA sequences for healthy individuals, these variants were predicted to be likely pathogenic and had been shown to be associated with Familial cancer of breast in ClinVar database. Through gene-gene interaction network, we found that MT-CYB gene was co-occurring significantly with other genes at Tibetan region. This had shed light on its possible involvement in hypoxia and low temperature adaptation. Cytochrome b protein is an integral membrane protein subunit of the cytochrome bc1 complex encoded by MT-CYB gene, this complex catalyzes the redox transfer of electrons from ubiquinone to cytochrome c in the mitochondrial electron transport chain. As the efficiency of the electron transport chain governs key aspects of aerobic energy metabolism [51], several investigators have suggested that functional modifications of redox proteins, such as cytochrome b, may be involved in physiological adaptation to different thermal environments [26, 27, 52]. It is interesting to note that variants located in CYB gene (such as C14766T) are known to be significantly higher in the high-altitude pulmonary edema a disorder arising due to acute exposure to high altitude above 3000□m and it has been hypothesized to play a role in high altitude sickness [46].

The formation of local co-occurrence pairs and similarity clustering divided the altitude groups into the higher and the lower altitude regions. This division might be possible due to the two reasons, (i) migration and demographic dynamics or (ii) process of selection on mitochondrial variants that in combination optimize mitochondrial bioenergetics in extreme conditions, experienced by these populations that lived at high altitude. Thus, the two node motifs identified at high altitude >3000m in the groups 7 and 8 can be suggested as candidate positions for a biological role in adaptation to these conditions. Overall, the co-occurrence network motifs provided detailed insight into finding the association of variable sites which are overlooked by haplogroup analysis alone and showed that selective pressures, such as high altitude, may generate constraints on the mitochondrial genomes, forcing the co-occurrence of certain variants on the mitochondrial genome.

## Methods

### Sequence Acquisition

We retrieved a total of 673 complete human mitochondrial DNA sequences of the healthy individuals from GenBank (http://www.ncbi.nih.gov/), where metadata was available (Supplementary file). These sequences were aligned using multiple sequence alignment tools, Clustal Omega [53] using default parameters. After alignment, all the sequences were mapped to master sequence, revised Cambridge Reference Sequence (rCRS). Since we were interested in altitude-wise stratification of these sequences based on available geographic information, we divided these sequences into eight altitude groups ranging from 0m to >4000m with an interval of 500m (Table 6) based on specific coordinates provided for each sequence in the published dataset (details provided in the Supplementary file). In this way we had eight altitude groups with different numbers of mtDNA sequences for construction of co-occurrence networks corresponding to each altitude group.

### Construction of Mitochondrial Motif and Gene-Gene Interaction Networks

**Step I**: For each altitude group we looked for any position within the samples having more than one allele and referred it as a variable site. For genomic equality, ambiguous nucleotides such as *X, M, Y*, etc were replaced with ‘N’ for all the sequences (Fig 1a). The sites with such nucleotides were not considered for constructing the network. These variable sites were then extracted from the aligned sequences for each group to further construct the co-occurrence networks or two node motifs as described in the step II.

**Step II**: Two variable sites are connected by an edge if co-occurrence frequency between these two does is greater than a given threshold value. The co-occurrence frequency *C*_*(i, j)*_ for a pair of nodes *i* and *j* is defined as,

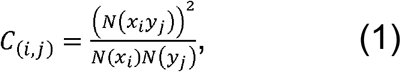

where *N(x*_*i*_*y*_*j*_) is number of times *x* and *y* nucleotides co-occur at *i*^th^ and j^th^ positions, respectively, *N(x*^*i*^) is total number of times nucleotide x occurs at i^th^ position and *N(y*_*j*_) is total number of times nucleotide y occurs at j^th^ positions.

The value of *C*_*(i,j)*_ ranges from 0 to 1, suggesting no co-occurrence (corresponding to 0) at all to a perfect co-occurrence between the i^th^ and j^th^ position. In such a scenario, two nodes are connected only if their co-occurrence frequency is equal to or greater than a given threshold value (C_th_). Putting C_th_ = 0 will give a globally connected network spanning all the nodes. Whereas, for C_th_ = 1, those nodes were connected which were perfectly co-occurring. We set the C_th_ to 1 and a co-occurrence motifs network for each sample was constructed for each altitude group (Fig 1b). Thus, for these co-occurrence networks, a node was represented by the position of variable sites and the edge between any two nodes was represented by co-occurrence frequency between those particular variable sites.

**Step III (Co-occurrence Network)**: From step II, we got as many co-occurrence networks as the number of samples in each group. The co-occurrence motif networks constructed for each sample in Step II for each group were merged to get a single master network for each altitude. Thus, the master motif network consists of all the motifs of that particular group (Fig 1c). In this way we constructed eight master networks, where nodes were variable sites and edges were co-occurrence frequency (Fig 9).

**Figure 9:** Co-occurrence network for Tibetan population (> 4000*m*) with co-occurrence frequency equal to 1. The colour is representing the degree of nodes. Blue for higher degree while green for intermediate degree and red for lower degree nodes. The figure was created using Gephi version 0.9.1 (https://gephi.org/) [60].

**Step IV (Gene-gene interaction Network)**: For each master network, the nodes (variable sites) were mapped to their corresponding genes to achieve one gene-gene interaction network for each altitude (Fig 1d). Since, two or more nodes may belong to the same gene or a gene pair, each link is counted as many times it is found in the co-occurrence network and this number was considered as weight for the corresponding gene-pair. For example, two-order motifs (3461-8715) and (4133-9157) belong to ND1-ATP6 gene-pairs, so this pair was counted twice for ND1-ATP6 gene pair, and so on. Finally, we have got two types of networks, first were master networks where nodes were variable sites and second were gene-gene interaction networks where nodes were genes.

### Classical Genetic Analysis and Machine Learning Classification of Altitude Groups

We perform the phylogenetic analysis to represent as the branching diagram the relationship between subjects. To reduce the influence from the population genetics, we first exclude those positions, which take part in the definitions of the haplogroups. Using the FastME algorithm we construct the phylogenetic tree and calculate distance matrix for all pairs of subjects [54]. Implementation of PCoA was taken from the Python package ‘Scikit-bio’ version 0.5.6 [55]. For the Random Forest algorithm, we treated variable mt-DNA sites as seed features to build and investigate binary classification models between each pair of altitude groups. The particular implementation was taken from Python package ‘Scikit-learn’ version 0.22; Python version 3.7.5 [56]. To perform classification between each pair of altitude groups we built a Random Forest model with the number of decision trees equal 500. 5-fold cross-validation was applied to test the effectiveness of a built model, yielding the average accuracy over cross-validated models. In DAPC (Discriminant Analysis of Principal Components), data is first transformed using a principal component analysis (PCA) and clusters are identified using discriminant analysis (DA). We apply it to each mitochondrial DNA region separately for 8 altitude groups of subjects. The DAPC algorithm is implemented in R package ‘adegenet’ version 2.1.3 [57, 58]. The fixation index (□□□□) is a measure of population differentiation due to genetic structure. We calculated the pairwise Fst test between all group pairs. Next, we performed 200 simulations, where each subject was randomly assigned to one of the altitude groups, Fst statistic was calculated for each simulation. We build a PDF for the calculated Fst statistic and compute an empirical p-value. The Fst calculating algorithm is implemented in R package ‘hierfstat’ version 0.5 [59].

### Null models

Random networks were generated for each sample of each altitude group. We took the same number of nodes as in the real network and a connection probability (*p*) based on the number of connections present in the real network, where p is defined as,

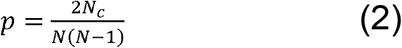

Here, Nc is the total number of connections and N is the total number of nodes in the corresponding real network. We compared the network properties of null networks with those of the real networks.

### Statistical analysis

A two proportion Z-test was used to find the significance in difference between the proportions of local connected nodes for a pair of the population i and j.

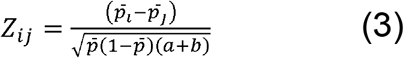

where, 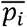 and 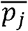 are proportions of the local connected nodes to total nodes for a pair of the altitude groups (*i* and *j*), 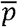 is the overall proportion calculated as a fraction of total local connected nodes to the total number of nodes for the two groups, a and b are inverse of the total number of variable sites of the altitude groups *i* and *j*, respectively. After calculating the z-score, we have decided the alpha value at 5%, for which the z-score is 1.96. A p-value is calculated using this Z-score. If p-value is less than 0.05, the difference in the proportions between the two altitude groups was considered to be significant.

### Similarity Clustering

We wanted to look for similarity between altitude groups. For this, we calculated Jaccard similarity coefficient using common nodes between each altitude group.

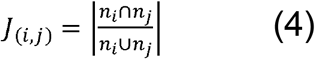

where, J_(i, j)_ is Jaccard similarity coefficient between two altitude groups *i* and *j*, and *n*_*i*_ and *n*_*j*_ are nodes of two altitude groups.

## Acknowledgments

SJ acknowledges support from SERB grant (EMR/2016/001921), the council of scientific and industrial research grant (CSIR, 25(0293)/18/EMR-II) Govt of India. SJ, MI and AK acknowledges the Ministry of education and science of the Russian Federation megagrant (075-15-2019-871). ADK acknowledges the council of scientific and industrial research grant (CSIR, 25(0293)/18/EMR-II) Govt of India for RA fellowship. P.S. thanks DST for the INSPIRE fellowship (IF150200). RKV gratefully acknowledges the University Grant Commission for SRF fellowship (305089).

